# Detection and isolation of H5N1 clade 2.3.4.4b high pathogenicity avian influenza virus from ticks (*Ornithodoros maritimus*) recovered from a naturally infected slender-billed gull (*Chroicocephalus genei*)

**DOI:** 10.1101/2025.11.28.689408

**Authors:** Dylan Andrieux, Manuela Crispo, Malorie Dirat, Laura Lebouteiller, Mathilda Walch, Emmanuel Liénard, Guillaume Croville, Maxime Fusade-Boyer, Loïc Palumbo, Julien Hirschinger, Guillaume Le Loc’h, Jean-Luc Guérin, Sébastien Soubies, Nicolas Gaide

## Abstract

Laridae birds, such as gulls, are known reservoirs of H13 and H16 low pathogenicity avian influenza virus (LPAIV) subtypes. However, during the recent outbreaks linked to the reemergence of high pathogenicity avian influenza virus (HPAIV) H5N1 clade 2.3.4.4b of the Goose/Guangdong lineage, European populations of Laridae birds suffered significant losses. HPAI cases were reported not only along the coastlines but also inland areas, particularly in France and Central Europe.

During a diagnostic investigation of a group of Laridae birds, part of a HPAIV outbreak reported in the South of France in 2023, larval stages of *Ornithodoros maritimus*, a nidicolous soft tick parasitizing seabirds, were recovered from a slender-billed gull (*Chroicocephalus genei*). Affected birds exhibited gross and histopathological lesions consistent with systemic HPAIV infection. Immunohistochemistry revealed marked neurotropism, oculotropism and multicentric epitheliotropism. Viral isolation and sequencing analysis confirmed the presence of HPAIV H5N1 clade 2.3.4.4b in both the gull and ectoparasites, showing from 99.64% to 100% nucleotide identity across five of eight RNA segments. While additional research is needed to properly assess the vector competence of *O. maritimus* for HPAIV, ticks may represent an interesting non-invasive surveillance tool for these viruses. This is the first time a HPAIV has been successfully isolated from tick larvae. These findings represent a first step toward understanding the potential role played by ticks in the spread of avian influenza viruses within marine bird colonies and among other ecosystems, considering the occurrence of specific behavioral traits, such as kleptoparasitim and the position of gulls at the interface between wild and domestic species.

## INTRODUCTION

High pathogenicity avian influenza (HPAI) is an infectious disease mainly affecting birds, caused by influenza A viruses (IAV) of the *Orthomyxoviridae* family. Since the end of 2020, high pathogenicity avian influenza virus (HPAIV) H5N1 clade 2.3.4.4b A/Goose/Guangdong/1/1996 lineage (Gs/Gd) has reemerged leading to seasonal outbreaks in a variety of wild and domestic birds (1). Aquatic birds, including Anseriformes (*i.e.* ducks, swans and geese) and Charadriiformes (*i.e.* shorebirds, terns and gulls), play an important role in the spread of avian influenza viruses (AIV) including HPAIV and represent well-established reservoir hosts (2).

Most gull species breed in large colonies (3) and are adapted to different environments, ranging from natural to urban areas, where they cohabit with wild waterfowl, poultry and humans (4). They are known for their kleptoparasitism, a foraging strategy consisting of “seizing food gathered by another individual” (5). These traits, together with their synanthropic and opportunistic behavior, increase the risk of HPAIV exposure in both wild and domestic birds, including poultry (6,7). While gulls are mostly known for being primary reservoirs of H13 and H16 low pathogenicity AIV (LPAIV) subtypes (8,9), their populations have experienced significant losses during the recent H5N1 outbreaks (10–14). In Europe, Laridae (*i.e.* gulls and terns) accounted for 17.4%, 51.4% and 20.6% of reported HPAI cases in free-living avifauna areas during the 2021-2022, 2022-2023 and 2023-2024 seasons, respectively (15–17). From a geographical point of view, the analysis of the distribution of cases between June 2022 and June 2023 showed that, in 2022, HPAI detections in Laridae were mainly located in the Channel, Atlantic and North Sea coastlines, while in 2023 many detections were observed along the coast but also inland areas, particularly in France and Central Europe (18). The overall number of HPAI cases and their evolving geographic distribution support the role of Laridae birds as key hosts in the current ecology of AIVs clade 2.3.4.4b H5Nx, at least for some of the circulating genotypes.

In addition, the recent literature highlights the potential of Laridae to carry and disseminate a variety of pathogens, including parasites, viruses and bacteria (19–21), emphasizing the complex interplay between wild birds and infectious agents. Specifically, the potential of carrying and disseminating parasites raises additional questions about other unknown or underestimated vectors in the transmission of AIVs and their ecological implications in both wildlife and peridomestic environments. Gulls carry numerous parasites including ectoparasites such as ticks (22), which are recognized vectors of several viral and bacterial agents (23–26). However, the potential involvement of ticks in AIV ecology needs to be further investigated. In particular, nidicolous ectoparasites, which inhabit host nests or burrows and feed on hosts intermittently, raise questions about their potential roles in virus persistence and environmental spread (27).

Here, we conducted a complete diagnostic investigation on five Laridae found dead during a HPAIV outbreak reported in the South of France in 2023. Among the five birds examined, soft ticks were recovered from one animal and further analyzed to assess the presence of HPAIV.

## MATERIALS AND METHODS

### Sampling

The five examined birds included three Mediterranean gulls (*Ichthyaetus melanocephalus*), one slender-billed gull (*Chroicocephalus genei*) and one sandwich tern (*Thalasseus sandvicensis*). All animals were found dead at Tartuguières, Grand Bastit, in the south of France, in 2023. One of the Mediterranean gulls (MG1) and the slender-billed gull (SB) were found on June 6^th^, 2023. The second Mediterranean gull (MG2) was collected on June 14^th^, 2023. The sandwich tern (ST) was found on July 5^th^, 2023, while the last Mediterranean gull (MG3) was discovered on July 12^th^, 2023. All carcasses, collected by the French national wildlife health surveillance network (SAGIR), had been frozen prior to laboratory submission for necropsy examination and diagnostic assessment.

### Necropsy

Complete necropsies were performed on thawed carcasses. Necropsy findings were recorded and lesions were documented. Cloacal and tracheal swabs (n=5) and, when present, immature growing feathers (MG1, MG2, MG3) were collected and stored at −80°C for molecular analysis. For histopathology and viral *in situ* detection, two birds, MG2 and SB, were selected based on *post-mortem* conditions. Specifically, tissue sections of brain, nasal cavity, trachea, lung, heart, liver, pancreas, gizzard, proventriculus, intestine, kidney, gonad, feathered skin, eye, uropygial, nasal and Harderian glands were collected and fixed in 10% neutral buffered formalin. Ticks identified on one bird (SB) were carefully removed and stored at −80°C in 1.5 mL Eppendorf tubes for molecular analysis.

### Parasitology

Ticks were morphologically examined using a stereomicroscope (Nikon SMZ, Tokyo, Japan) and identified based on standard morphological keys (28–30).

### Histopathology

Upon formalin fixation, tissue sections and whole ticks were routinely paraffin-embedded and sectioned at 3 μm. Slides were stained with hematoxylin and eosin (H&E) and examined by light microscopy and scanned at 20× magnification using VS200 Olympus slide scanner.

### Immunohistochemistry

For viral *in situ* detection and tissue distribution, serial sections, 3 μm-thick, were mounted on positively charged slides and stained with immunohistochemistry (IHC). The immunohistochemical assay was performed using a monoclonal mouse antibody directed against influenza A virus (IAV) nucleoprotein (NP) (Clone H16-L10-4R5 HB-65, reference BXC-BE0159, Biozol, Germany) (31). Sections of IAV positive tissues were used as positive controls. Negative controls included sections incubated without the primary antibody or with a monoclonal antibody belonging to the same isotype (IgG2) while sections of AIV positive tissues were used as positive controls. 3,3’-Diaminobenzidine (DAB) was used for chromogenic reaction and antigen detection, except for tegument and ocular tissues for which magenta chromogen (ENVISIO FLEX HRP Magenta system; Agilent, USA) was used to avoid misinterpretation with endogenous pigments. Parenchymal immunoreactivity of each organ was semi-quantitatively scored according to distribution (32). Slides were additionally scanned at 20× magnification using a VS200 Olympus slide scanner.

### Virology

In order to assess the presence of AIV on the surface and/or within the ticks collected from the slender-billed gull and to assess their infection status, viral isolation was attempted. The ectoparasites were placed in 1 mL of phosphate-buffered saline (PBS) containing 0.2% (m/v) bovine serum albumin, 10,000 UI/mL penicillin, streptomycin and amphotericin B (hereafter referred to as “diluent”) and incubated for 5 minutes at room temperature. The diluent was then removed and used as the “outside” sample. Subsequently, ticks were cut into pieces using a sterile scalpel blade, transferred into 1 mL of fresh diluent and mechanically disrupted using metal beads under agitation. The homogenate, representing the “inside” sample, was then inoculated into 10-day-old specific pathogen-free (SPF) embryonated chicken eggs (PFIE INRAe, Nouzilly, France), after being serially diluted from 10^-1^ to 10^-4^. Six eggs were inoculated with 100 µL of dilution into the allantoic cavity for each condition (outside and inside samples) and dilution. Two days post-inoculation, allantoic fluids were collected and hemagglutination assays (HAA) were performed as previously described (31). Viral titers, expressed as median egg infectious doses / mL (EID_50_/mL), were calculated using the “midSIN” calculator (33).

### Molecular biology

For each bird, tracheal and cloacal swabs were pooled together. Subsequently, upon nucleic acid extraction, AIV H5 RT-qPCR (34) was performed on tracheal and cloacal swab pools and available immature feathers using the Qiagen kit QuantiNova probe (Qiagen, Hilden, Germany), as previously described (35). RT-qPCR on the brain, uropygial gland and tracheal swab collected from the SB was performed using the IDvet Influenza A Triplex kit (IDvet, France).

### Nanopore sequencing

Nanopore sequencing was done as previously described by Croville *et al.* (36). RNA extraction of the gull tracheal swab, brain and uropygial gland as well as the allantoic fluid containing the tick virus was performed using MagFAST Extraction Kit with IDEAL32 according to the manufacturer’s instructions. Reverse-transcription was done with Uni-12 primers (37) and RevertAid cDNA Synthesis Kit (ThermoFisher, Waltham, USA). Then, PCR was done with Phusion hot start flex kit (ThermoFisher) and Uni12/Uni13-based primers (38). After electrophoresis on a 1% gel, bands at the expected size were cut and purified with MN Nucleospin Gel and PCR clean-up kit (Macherey-Nagel), following manufacturer’s instructions.

For the tick viral isolate obtained from embryonated chicken eggs (“inside” sample), nanopore sequencing of PCR amplicons was performed by Eurofins genomics (Ebersberg, Germany).

For the bird (SB) visibly parasitized with ticks, PCR products of each selected matrix (brain, uropygial gland and tracheal swab) were pooled and purified with AMPure XP beadx. The DNA library was prepared using the SQK-NBD114.24 sequencing kit (Oxford Nanopore Technologies - ONT, Oxford, United Kingdom). The DNA library was loaded on a R10.4.1 MinION flow cell and the sequencing run was launched on a MinION Mk1C device (ONT). High accuracy base-calling was performed in real-time with Guppy (v3.5) embedded in the MK1C software (v19.12.12).

### Sequencing data analysis

The FASTQ files were aligned using minimap2 (39) and the SAMtools (40) package. The whole-genome consensus sequences were generated using the consensus command on the iVar (41) pipeline that embeds the SAMtools mpileup tool (42).

The viral sequences obtained during this study are available on GenBank, accession numbers from PZ225337 to PZ225365.

### Phylogenetic analysis

Phylogenetic analysis was performed on the segment encoding hemagglutinin (HA), the preferred target for accurately tracking the evolution of influenza A viruses and for clade classification, as well as on the other viral segments that could be sequenced in all samples (polymerase basic 2 (PB2), NP, matrix (M) and non-structural (NS) segments) for genotype identification. Consensus sequences of each viral gene segment, obtained from the egg-passaged, tick-derived virus and the three different matrices from the SB, were supplemented with closely related sequences identified via BLAST on GISAID (https://www.gisaid.org/), along with reference sequences of HPAI clade 2.3.4.4b H5Nx corresponding to well-characterized European genotypes (43). These sequences were selected because they define the genetic diversity and genotypic classification within this clade. Multiple sequence alignment was then performed using MAFFT version 7 (https://mafft.cbrc.jp/alignment/server/index.html). IQ-TREE version 1.6.12 (http://www.iqtree.org/) was then used to select the most suitable model for each gene segment based on the Bayesian Information Criterion (BIC) and to perform maximum likelihood phylogenetic analyses with 100,000 ultrafast bootstrap replicates for each segment. Phylogenetic trees were then visualized using FigTree version 1.4.2. (http://tree.bio.ed.ac.uk/ software/figtree/).

## RESULTS

### Gross lesions

Two animals (ST, MG3) exhibited extensive autolytic changes and advanced decomposition, preventing proper assessment of physiological traits, such as gender, and the identification of significant lesions. The rest of the birds examined included two adult, sexually inactive females (MG1, MG2) and one adult, sexually inactive male (SB).

Upon external examination, numerous ectoparasites were discovered firmly attached to the skin of the vent and the ventral surface of the left cervico-thoracic region of the SB. The ectoparasites were round to oval in shape, with a smooth surface, black or dark red in color, and ranged between 2 and 5 mm in size (Fig.1A). Small to moderate numbers of oval, cream-colored structures, 1 to 2 mm in size, were also identified in the subcutis and fascia of adipose tissue of the cervico-thoracic region, bilaterally in the same bird.

**Figure 1.**
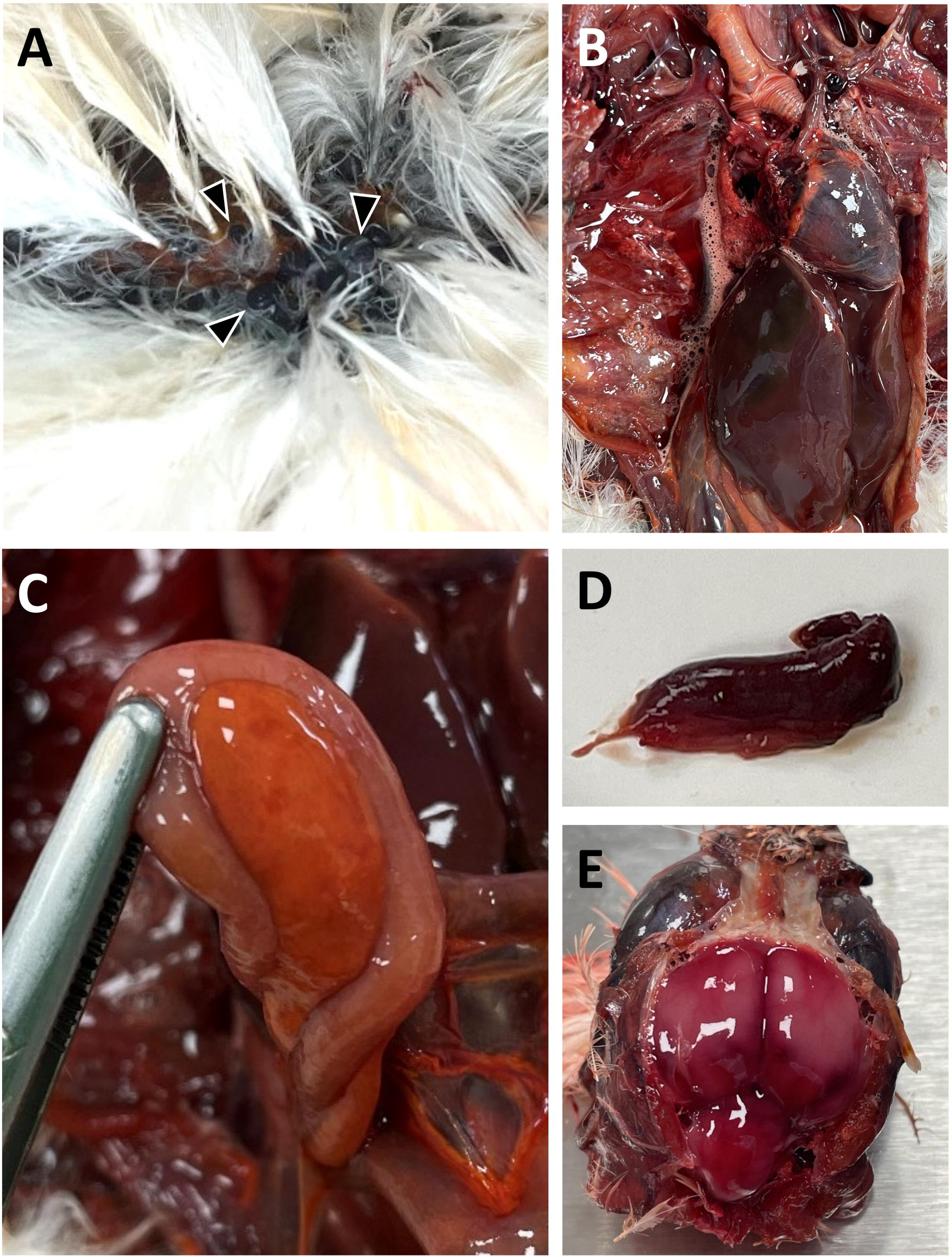
Gross findings of a slender-billed gull naturally infected with H5 highly pathogenic avian influenza viruses (HPAIV) (A) Numerous ticks were parasitizing the dead gull. (B) Heart, lungs and liver appear congested and edematous. (C) Pancreas is mottled and pale. (D) Splenomegaly. (E) Brain is diffusely red.

The majority of the organs, particularly the heart, liver and lungs, appeared diffusely and markedly congested and edematous (MG1, SB) with concurrent pleural and coelomic serous effusions (SB) (Fig.1B). The pancreas was moderately mottled, diffusely pale or yellow-orange in color (MG1, SB) (Fig.1C). Mild to moderate splenomegaly was observed in two birds (MG1, SB) (Fig.1D). The brain appeared diffusely red (Fig.1E). Scant amounts of ingesta were present in the proventriculus and gizzard. The latter showed diffuse mucosal thickening and a bile-stained, friable koilin. Kidneys were moderately mottled, with an increased lobular pattern, and pale striations (MG1, SB).

### Histopathology

The vast majority of tissues examined showed marked generalized congestion and moderate freezing artefacts (Fig.2A). Necrotic-inflammatory changes predominated in both selected birds (MG2 and SB). Specifically, the pancreas revealed severe, multifocal to coalescing necrosis of acinar cells (Fig.2B) with concurrent marked vasculitis (fibrinoid necrosis) and mesenteric steatonecrosis. Segmental epithelial necrosis and intrafollicular debris were observed in the growing feather follicle (Fig.2C). The uropygial gland showed mild, multifocal segmental lobular necrosis (Fig.2D). In MG2, mild cerebro-cortical neuronal degeneration and necrosis, with multifocal glial nodules, were noticed in the brain. Ocular sections exhibited moderate to marked multifocal, bilateral, lympho-plasmacytic infiltration of the ciliary body and irido-corneal angle, while the retina was within normal limits (Fig.2E). Moderate multifocal necrosis of the adrenal parenchyma was also present.

**Figure 2.**
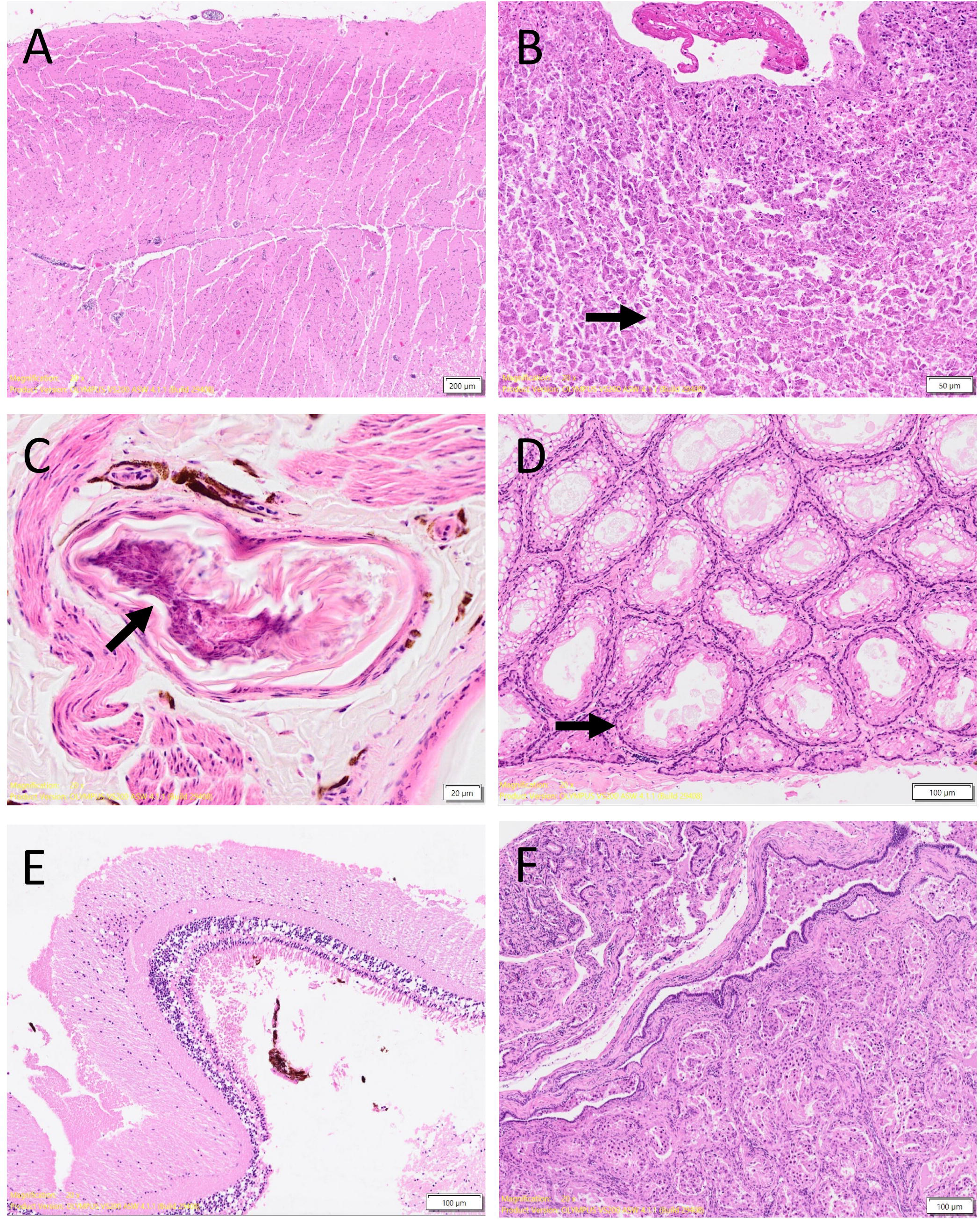
Histopathological findings in tissues from slender-billed gulls naturally infected with H5 highly pathogenic influenza virus (HPAIV). (A) Brain, (B) pancreas, (C) growing feather, (D) uropygial gland, (E) retina, (F) testis and epididymis. Most tissues are within normal histological limits, with frozen and autolytic artefacts. Notable lesions include peripancreatic fibrinous exudate (B, arrow), segmental epithelial necrosis and intraluminal debris in the growing feather follicle (C), uropygial gland parenchyma (D), and epididymal lumen (F).

In the SB, rare foci of myocardial degeneration and necrosis, associated with minimal lymphocytic infiltration, were present in the right ventricle. Skeletal muscles revealed marked, focally extensive degeneration and necrosis of myofibers, with concurrent lymphohistocytic infiltration. Testicles presented moderate amounts of sloughed epithelial cells and cell debris within the lumen of tubules, rete testis and epididymis, with few foci of tubular degeneration, and spermatozoa rarefaction (Fig.2F). The Harderian gland exhibited mild, multifocal parenchymal necrosis.

Incidental findings included: small numbers of deutonymphs (hypopi) of hypoderatid mites (*Hypodectes* spp.) scattered in the subcutis of the cervico-thoracic region with minimal inflammatory response (SB); mild esophageal capillariosis (SB); mild fibrino-heterophilic proventriculitis and ventriculitis with rare intralesional bacterial colonies (SB); mild duodenal cestodiasis (SB); mild proventricular nematodiasis (SB); nonspecific, multifocal accumulation of brown-black, granular material within the walls of tertiary bronchi, Kupffer cells and splenic phagocytic cells (SB and MG2); mild multifocal foci of renal tubular mineralization (SB), tubular degeneration and necrosis (MG2).

Overall, multi-centric visceral necrotizing and nonsuppurative inflammation was consistent with a systemic infection (e.g. viral etiology), associated with pancreatitis, encephalitis, myocarditis/myositis, endophthalmitis and adenitis, and concurrent polyparasitism for the slender-billed gull.

### Immunohistochemistry

Viral antigens (NP) were detected systemically in both selected birds (Table 1). The most affected organs, in terms of frequency and/or intensity of detection, included: brain, eye, pancreas, feathered skin, uropygial and nasal glands (Fig.3). Viral antigens were identified within necrotic-inflammatory areas and a variety of cells (nuclear and cytoplasmic positivity) (Table 1).

**Table 1.** Viral antigen distribution in different organs of the slender-billed gull (SB) and Mediterranean gull (MG2). For each bird and each tissue, viral antigen detection is expressed as a score according to Landmann *et al.* (2021): 0 (non), 1 (focal/oligofocal), 2 (multifocal), 3 (coalescing/widespread). NA: not available.

|  |  |  |  |  |
| --- | --- | --- | --- | --- |
| **Nervous** | Brain | 3 | 2 | Neurons, Purkinje cells (cerebellum), glial cells, ependymal cells. |
|  | Eye | 2 | 1 | Iridocorneal angle (*trabeculum*), iris, ciliary body, retina, optic nerve, pecten |
| **Respiratory** | Lung | 0 | 0 | - |
|  | Nasal cavity | 0 | 0 | - |
|  | Trachea | 0 | 0 | - |
| **Cardiovascular** | Heart | 2 | 0 | Cardiomyocytes |
| **Digestive** | Esophagus | 0 | NA | - |
|  | Pancreas | 2 | 2 | Exocrine acinar cells |
|  | Intestine | 1 | 0 | Myenteric plexus, mucosal epithelium (autolysis) |
|  | Liver | 0 | 0 | - |
|  | Gizzard | 0 | 0 | - |
|  | Proventriculus | 1 | 0 | Glandular epithelial cells |
| **Urinary** | Kidney | 2 | 1 | Renal tubular epithelium |
| **Tegumentary** | Feathered skin | 2 | 1 | Epithelial cells of growing feathers, epithelial lining of the dermic papilla of resting feathers |
| **Endocrinian** | Uropygial gland | 2 | 2 | Glandular epithelial cells |
|  | Nasal gland | 2 | 2 | Glandular epithelial cells, luminal debris |
|  | Gonad | 2 | 0 | Seminiferous tubules, epithelium of epididymis and luminal content |
|  | Adrenal gland | 3 | 0 | Glandular epithelial cells, interstitium |
|  | Harderian gland | 0 | NA | - |
| **Locomotor** | Skeletal muscle | 1 | 1 | Myofibers |

**Figure 3.**
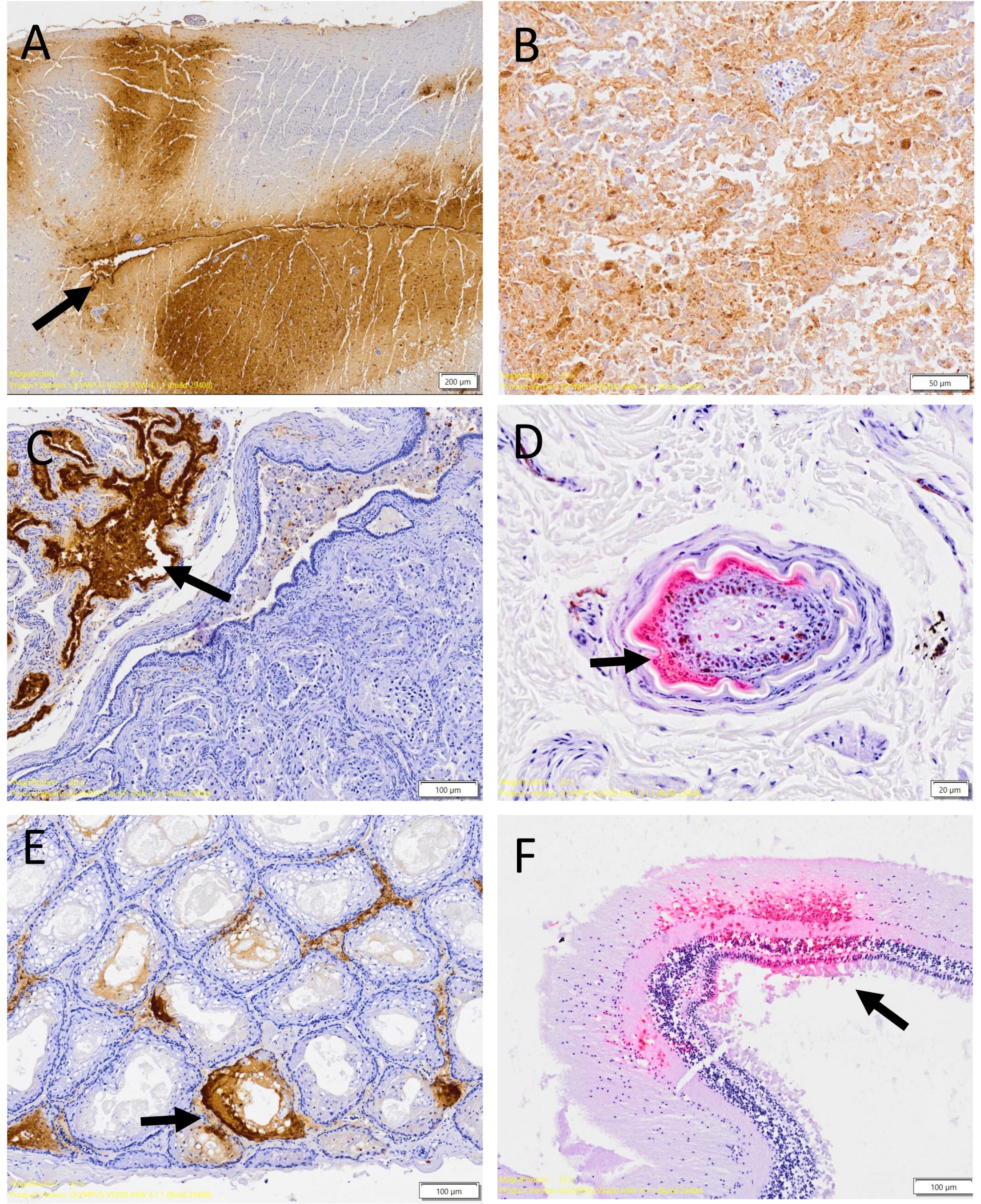
Immunohistochemical detection of influenza A nucleoprotein (NP) in tissues from slender-billed gull naturally infected with H5 highly pathogenic avian influenza virus (HPAIV). (A) Brain, (B) pancreas, (C), testis and epididymis, (D) growing feather, (E) uropygial gland, (F) retina. For eye and skin samples (D and F), a magenta chromogenic substrate was used to avoid interference from endogenous brown/black pigments; for other organs, DAB (brown color) was used.

### Molecular biology

All cloacal and tracheal swab pools (5/5) and immature feathers (3/3) were positive for H5 AIV by RT-qPCR (Table 2). The mean Cq for the tracheal and cloacal swab pools was higher (30.71) than for immature feathers (23.77) (Table 2).

**Table 2.**
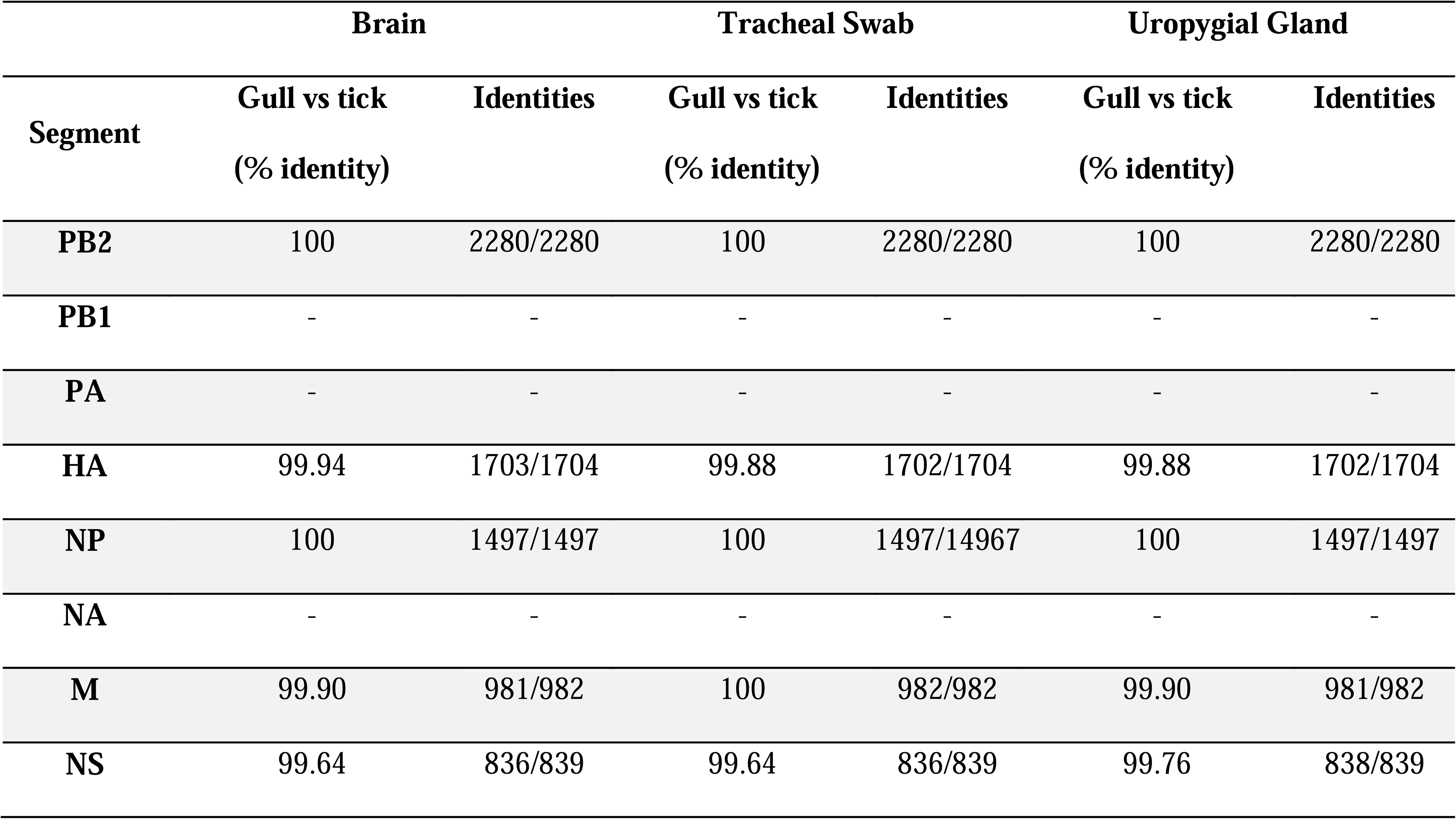
RT-qPCR Cq of tracheal and cloacal swabs pools and immature feathers of the five birds naturally infected with an H5N1 influenza virus. . For each bird, tracheal and cloacal swabs were collected and pooled before being analyzed by RT-qPCR to detect H5 AIV. Immature feathers were analyzed when found on the bird. NA = not available. Detection threshold: Cq ≤ 38.

In addition, tracheal swab, brain and uropygial gland of the slender-billed gull showed Cq values of 17.95, 12.68 and 23.65 respectively.

### Parasitology

Only soft tick larvae, unengorged and engorged, were recovered from the SB (Fig.4A). All specimens were identified as *Ornithodoros maritimus* (44) based on the presence of morphological features included in the *Ornithodoros capensis sensu lato* complex, such as the apical dentition of the subparallel hypostome with a bluntly rounded apex (Fig.4 B-D).

**Figure 4.**
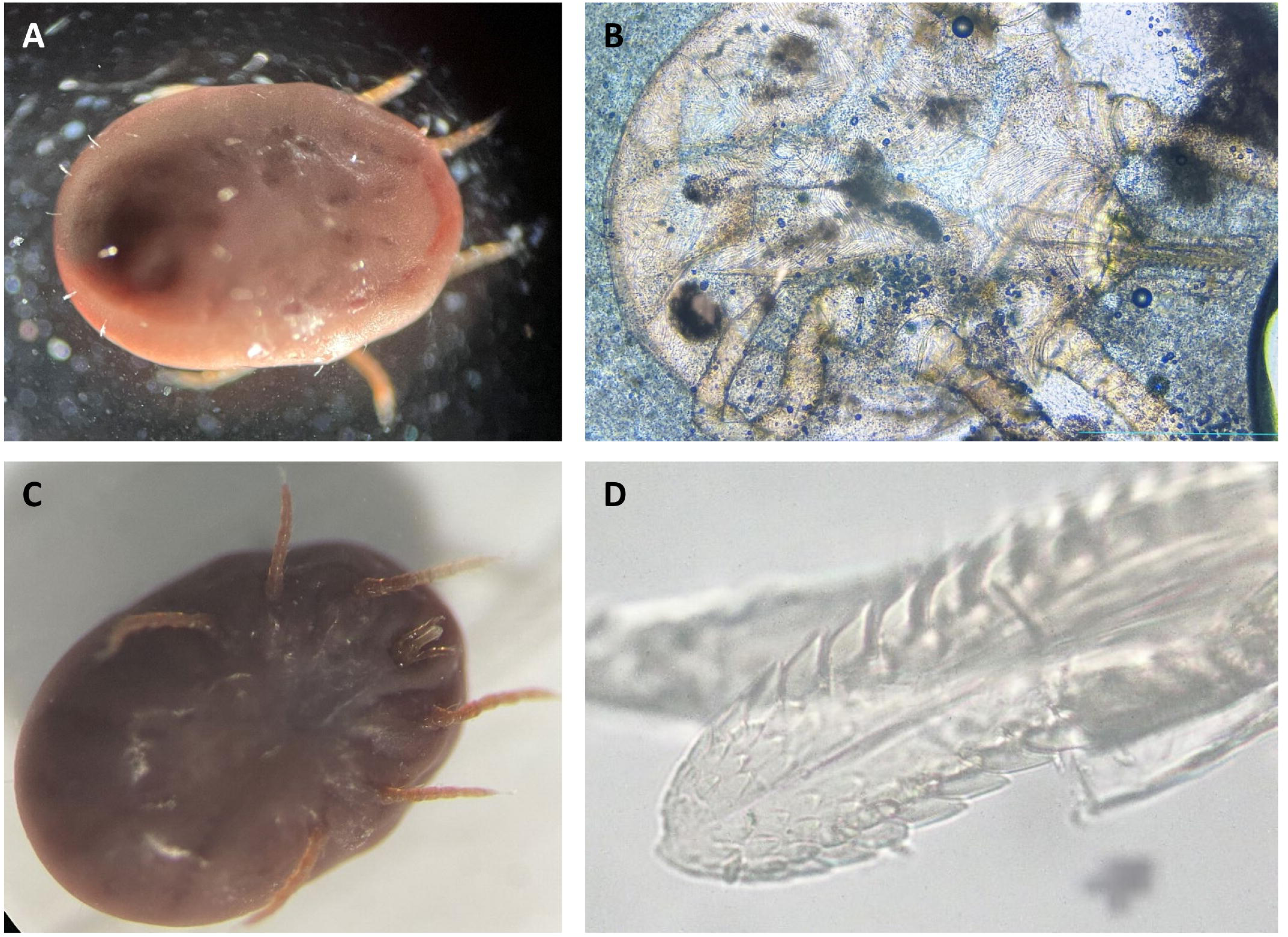
Tick morphological identification. (A) Unengorged tick larva (3 pairs of legs), dorsal view. (B) Tick larva observed in wet mount. (C) Unengorged tick larva, ventral view. (D) Tick subparallel hypostome observed under an optical microscope.

### Viral isolation

The allantoic fluids collected from the SPF eggs inoculated with the “outside” sample all tested negative by HAA (6/6, no mortality). As a result, the titer of the “outside” sample was estimated to be < 10^0.777^ (EID_50_/mL).

The allantoic fluid obtained from the SPF eggs inoculated with the “inside” sample, corresponding to the shredded ticks, was positive by HAA. Specifically, all SPF eggs inoculated with the 10^0^ and 10^-1^ dilutions of the “inside” sample died 48 hours post-infection and their allantoic fluid was positive by HAA. The allantoic fluid collected from 4/6 eggs inoculated with the 10^-2^ dilution was also positive. Based on these results, the initial titer of the “inside” sample was estimated to be 10^2.963^ [2.496;3.343] (EID_50_/mL) [+/- 95% credible interval].

### Sequencing and phylogenetic analysis

Gull-associated viral sequences from the tracheal swab, brain and uropygial gland of SB, and the tick virus isolated from the “inside” sample, were determined and compared. Five of eight influenza RNA segments (PB2, HA, NP, M and NS) showed 99.64% to 100% identity (Table 3). Unfortunately, PB1, polymerase acidic (PA) and neuraminidase (NA) segments of the tick virus did not yield sequences of sufficient quality for identity or phylogenetic analysis.

**Table 3.**
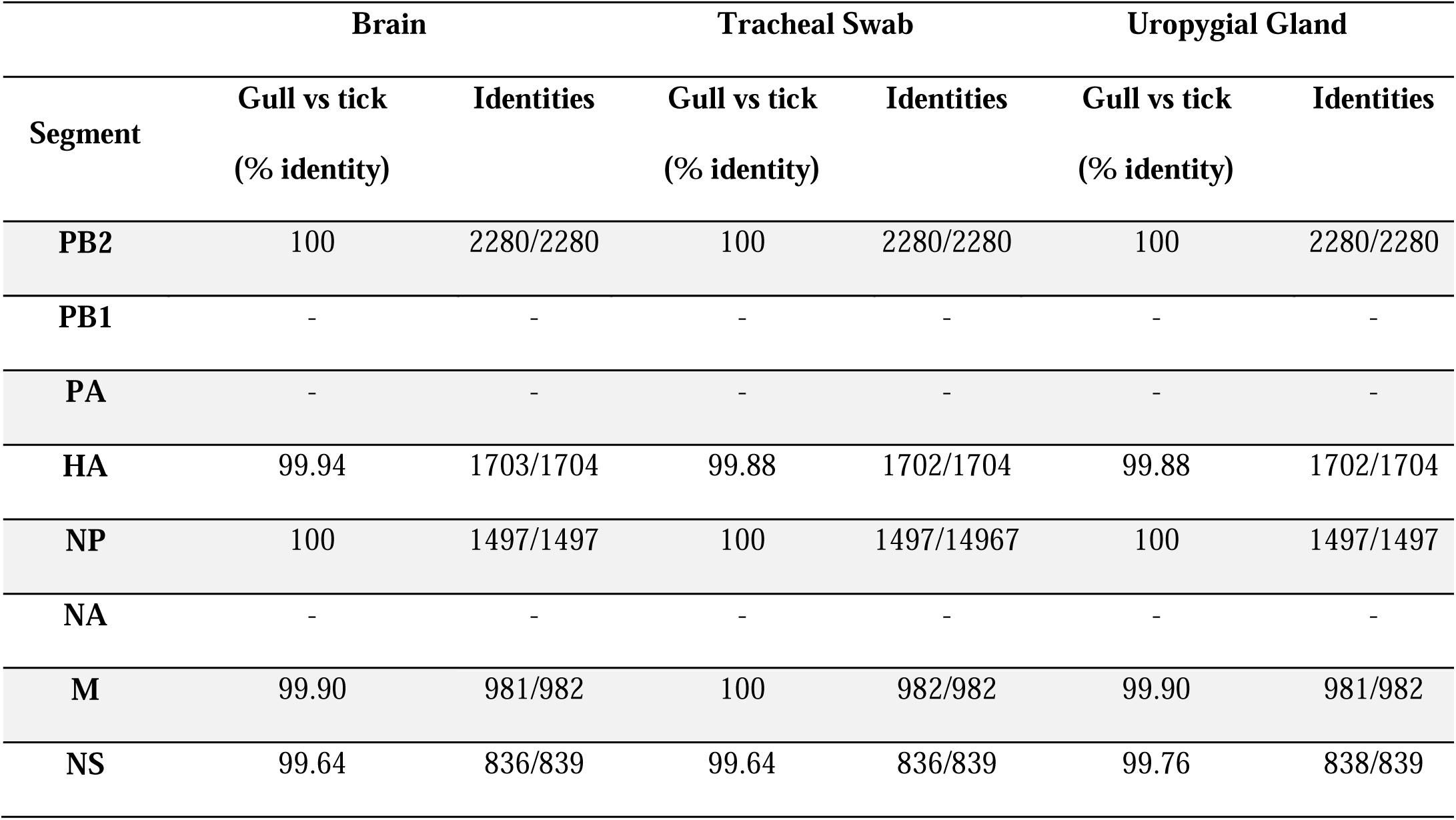
Percentage of identity between gull and tick viruses. The gull virus was extracted from the brain, tracheal swab and uropygial gland, the tick virus was amplified in chicken embryonated eggs before extraction from the allantoic fluid. PB1 could not be sequenced for the tick virus, PA and NA were not analyzed due to poor quality of the tick virus sequences for these segments.

Sequence analysis showed 1 or 2 nucleotide substitutions in the HA segment of the tick virus compared to the gull virus, depending on the avian sample considered, which did not result in amino acid changes.

Phylogenetic analyses on the five viral segments available from both the gull and the tick showed that their sequences clustered together, which was also supported by ultrafast bootstrap values, suggesting close similarity (Fig.5 and Supplemental Figures 1 to 4). Bootstrap support was high across all segments (90-96%), with the strongest values observed for HA (96%) and NS (95%), while PB2, NP and M showed slightly lower but still relatively high support (90%, 93% and 92%, respectively). This variability in bootstrap support likely reflects differences in the level of genetic diversity across segments. For PB2 and M in particular, the sequences obtained from the gull and the tick were identical to each other, but also identical to several closely related reference sequences, resulting in a limited number of phylogenetically informative sites and consequently slightly lower bootstrap values. The three other viral segments, only available from the gull, also clustered together (Supplemental Figures 5 to 7). This suggests the presence of a single viral lineage in both the bird and the parasite. However, due to the lack of complete genome sequences, the genotype of these viruses could not be definitively determined, although phylogenetic analysis suggests that both viruses belong to either the BB or DF genotype.

**Figure 5.**
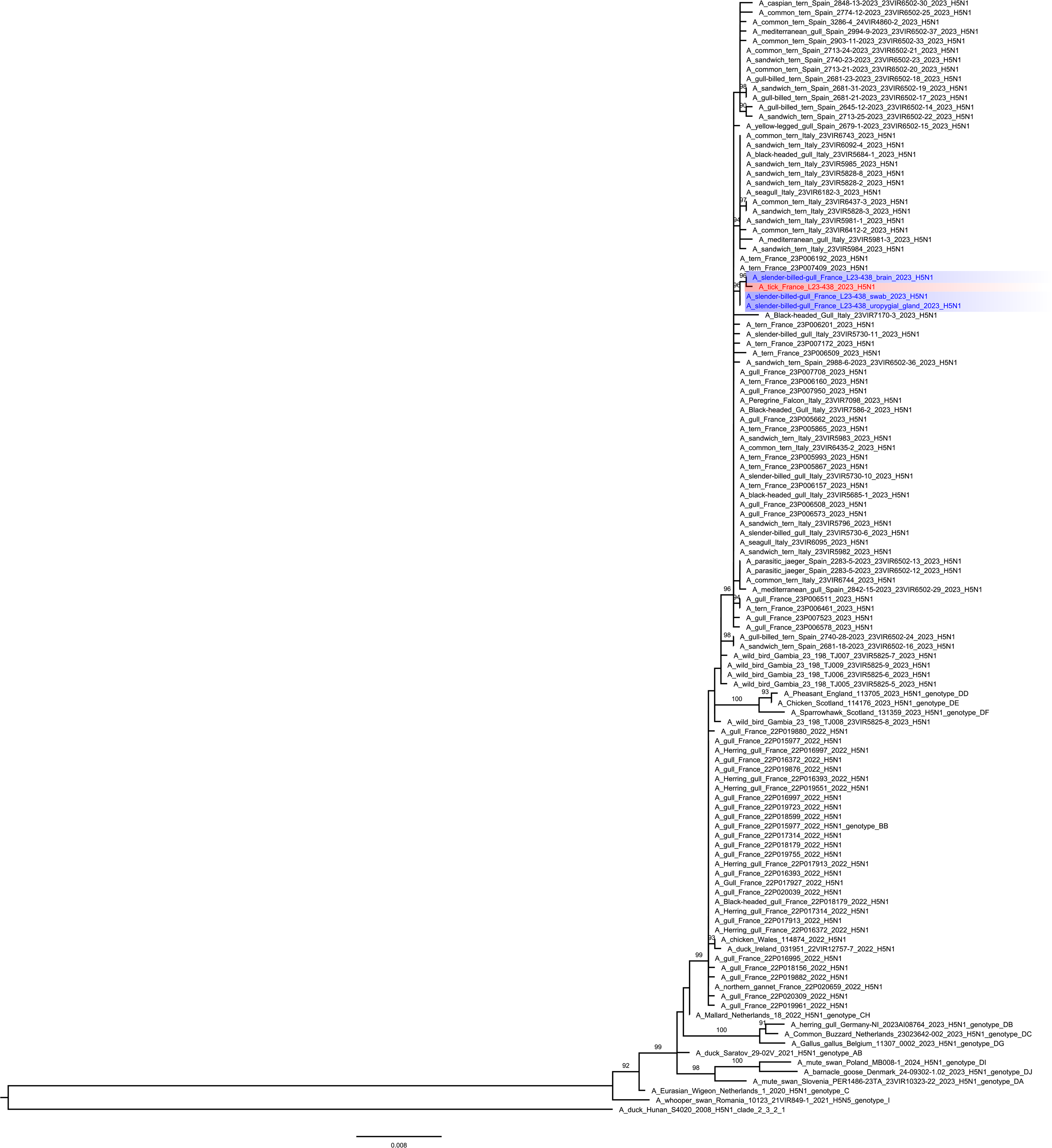
Maximum-likelihood phylogenetic tree of the hemagglutinin (HA) gene showing the clade 2.3.4.4b H5N1 virus isolated from a tick collected from a slender-billed gull, alongside the virus detected in the same gull. The tree includes the closest related strains as well as reference sequences representing European genotypes, all belonging to clade 2.3.4.4b. The virus detected in the tick is shown in red, while the virus detected in the gull is shown in blue. The tree was rooted using strain A/duck/Hunan/S4020/2008 (H5N1, clade 2.3.2.1). Ultrafast bootstrap support higher than 90% is indicated above the nodes. The scale bar indicates the number of nucleotide substitutions per site.

## DISCUSSION

In this study, we detailed the pathological presentation and viral antigen tissue distribution of H5N1 clade 2.3.4.4b HPAI in naturally-infected wild gulls, exhibiting marked neurotropism, oculotropism and multicentric epitheliotropism over the course of a systemic infection. We were also able to successfully detect the virus in ectoparasites, specifically tick larvae identified as *Ornithodoros maritimus* recovered from one of the infected birds, a slender-billed gull. *Ornithodoros maritimus* is a soft tick (Argasidae) commonly reported in a wide range of seabirds, including gulls (44,30). Our findings provide a first step towards a better understanding of the role played by ticks in the diffusion of AIVs among marine birds. In this context, some studies have focused on other arthropods, such as blowflies (genus *Calliphora*) and mites (*Dermanyssus gallinae*), which have been identified as potential vectors for AIV in Japan and Germany, respectively (45–47). To the best of our knowledge, this is the first report of HPAIV detection, both by molecular analysis and viral isolation, in ticks visibly parasitizing birds. Nevertheless, there are still several questions to address.

Viral genome sequences obtained from the tick and the gull showed very high nucleotide identity, ranging from 99.64% to 100% across five of eight segments, depending on the avian matrix tested (tracheal swab, brain, and uropygial gland). There was no significant difference between the viruses derived from the three avian matrices selected. This finding was further supported by phylogenetic analyses, in which viral segments from the tick and gull samples consistently clustered together. Bootstrap support was high across segments (90-96%), with HA and NS showing the stronger values, followed by NP, PB2 and M. These results are consistent with the circulation of a closely related viral lineage shared between the two hosts.

The soft tick *Ornithodoros maritimus* belongs to the Argasidae family and its ecology is highly adapted to seabirds. It is predominantly distributed in the Western Palearctic, where its role as a parasite of seabirds such as the Mediterranean storm petrel (*Hydrobates pelagicus melitensis*) has been extensively documented (30,48,49).

Unlike hard ticks, this species feeds multiple times during its nymph and adult stages, increasing the risk of viral acquisition and transmission within a bird colony (50). In our study, only larvae were sampled. Since larvae feed only once, with a blood meal lasting 4 to 6 days under experimental conditions (51), our results may reflect the accidental capture of a tick feeding on an infected bird. This observation raises the question of the virus’s ability for transstadial persistence to later developmental stages. The epidemiological consequences would differ if the nymphal and adult stages were also involved, as a new blood meal on a host visiting the same nest could result in horizontal transmission. However, the nidicolous lifestyle of *O. maritimus* may limit viral spread between bird colonies, since ticks tend to remain on hosts within the same nest. In addition, gulls exhibit a strong site fidelity to their nesting areas (50,52,53).

Additional research is needed to properly assess the vector competence of *O. maritimus*, both mechanically and biologically. On the one hand, mechanical transmission may result from external contamination of the tick’s rostrum or through allopreening, which can facilitate the removal of ectoparasites. On the other hand, biological transmission, which is not supported by current literature on avian influenza or our data, would require efficient viral replication within the tick. In both scenarios, viraemia would be a major prerequisite, as the initially infected host would need to have circulating virions in the bloodstream in order for the tick to become infected. Addressing these questions would require additional studies, which are beyond the scope of this report.

Our results show that soft ticks may represent an environmental sample of interest for HPAIV surveillance. Harvesting engorged soft ticks in seabird nests may be a suitable and practical method to assess the circulation of HPAIV in a bird colony. The presence of HPAIV in other tick species should also be explored to understand if their collection and testing could be used as an additional and non-invasive surveillance tool for influenza in wild birds.

Gulls are a reservoir for HPAIV, and are located at the interface between wild and domestic avian species. Breeding behaviors, such as nest fidelity and chick brooding could expose birds to nidicolous ticks repeatedly, potentially facilitating HPAIV circulation within a colony. Considering the proximity of gulls to other wild birds, poultry farms and mammals (54), these findings raise important questions about the role of ectoparasites, particularly soft ticks, in the transmission of HPAIV. For example, in Finland, HPAI-related morbidity and mortality events were reported in farmed red and silver foxes (*Vulpes vulpes*, *Vulpes lagopus*), minks (*Neovison vison*) and racoon dogs (*Nyctereutes procyonoides*) and ultimately linked to mass mortality of black-headed gulls (*Chroicocephalus ridibundus*). Gulls have been observed regularly visiting several lakes in the area and feeding on the farms (54).

Interestingly, and as previously reported by several research groups, immature feathers are the most suitable matrix to detect HPAIV infection in clinically affected birds compared to swabs (55–58). Indeed, the RT-qPCR Cq values were significantly lower for immature feathers (mean Cq: 23.77) than for the pool of tracheal and cloacal swabs (mean Cq: 30.71) Our results support the fact that immature feathers, when available, should be routinely sampled for HPAIV surveillance in wild as well as domestic birds.

The detection of a H5N1 HPAIV in *O. maritimus* ticks and an infected gull underscores the complex interplay between biological vectors, host behavior, and environmental factors. A better understanding of these dynamics is essential for assessing the risk of viral spillover into domestic avian populations and for improving surveillance strategies in seabird colonies.

## Abbreviations

AIV: avian influenza viruses
BLAST: basic local alignment search tool
HPAIV: high pathogenicity avian influenza virus
HA: hemagglutinin
HAA: hemagglutination assay
IHC: immunohistochemistry
IAV: influenza A virus
LPAIV: low pathogenicity avian influenza virus
MG: Mediterranean gull
M: matrix
NA: neuraminidase
NP: nucleoprotein
NS: non-structural
PA: polymerase acidic
PB: polymerase basic
PBS: phosphate-buffered saline
RT-qPCR: quantitative reverse transcription polymerase chain reaction
SB: slender-billed gull
SPF: specific pathogen-free
ST: sandwich tern.

## ACKNOWLEDGMENTS

The authors acknowledge the « Chaire de Biosécurité et Santé Aviaires », hosted by the National Veterinary School of Toulouse (ENVT) and funded by the Direction Générale de l’Alimentation, Ministère de l’Agriculture et de la Souveraineté Alimentaire, France; as well as the “Chaire de Professeur Junior” held by Sébastien Soubies and funded by “Agence nationale de la recherche” and by the Department of Animal Health of the Institut national de recherche pour l’agriculture, l’alimentation et l’environnement (INRAe). We sincerely thank Kathleen Perrot from the site of Tartuguière for her valuable collaboration in this study.

We gratefully acknowledge the authors, originating and submitting laboratories of the sequences from GISAID on which this research is based. All submitters can be contacted via www.gisaid.org.

---

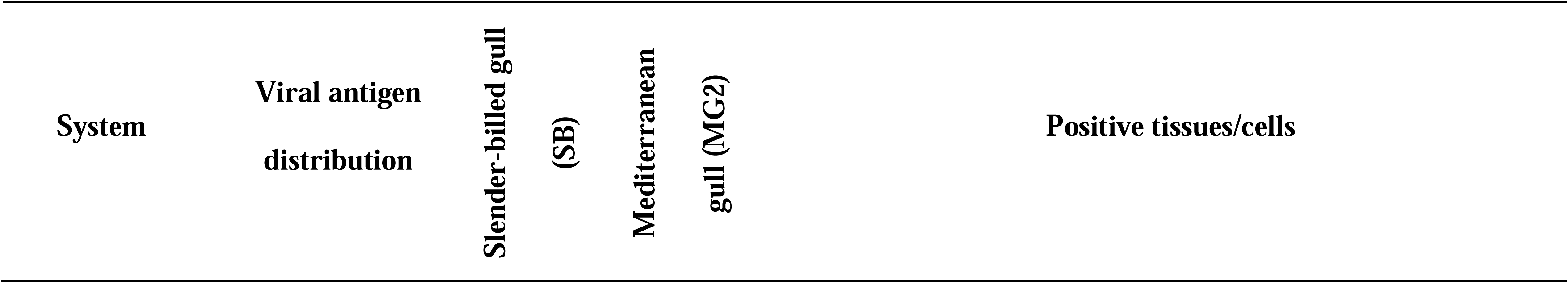

**Supplementary figure 1.** Maximum-likelihood phylogenetic tree of the PB2 gene showing the clade 2.3.4.4b H5N1 virus isolated from a tick collected from the slender-billed gull, alongside the virus detected in the same gull. This unrooted tree includes the closest related strains as well as reference sequences representing European genotypes, all belonging to clade 2.3.4.4b. The virus detected in the tick is shown in red, while the virus detected in the gull is shown in blue. Ultrafast bootstrap support higher than 90% is indicated above the nodes. The scale bar indicates the number of nucleotide substitutions per site.

**Supplementary figure 2.** Maximum-likelihood phylogenetic tree of the NP segment showing the clade 2.3.4.4b H5N1 virus isolated from a tick collected from the slender-billed gull, alongside the virus detected in the same gull. This unrooted tree includes the closest related strains as well as reference sequences representing European genotypes, all belonging to clade 2.3.4.4b. The virus detected in the tick is shown in red, while the virus detected in the gull is shown in blue. Ultrafast bootstrap support higher than 90% is indicated above the nodes. The scale bar indicates the number of nucleotide substitutions per site.

**Supplementary figure 3.** Maximum-likelihood phylogenetic tree of the M gene segment showing the clade 2.3.4.4b H5N1 virus isolated from a tick collected from the slender-billed gull, alongside the virus detected in the same gull. This unrooted tree includes the closest related strains as well as reference sequences representing European genotypes, all belonging to clade 2.3.4.4b. The virus detected in the tick is shown in red, while the virus detected in the gull is shown in blue. Ultrafast bootstrap support higher than 90% is indicated above the nodes. The scale bar indicates the number of nucleotide substitutions per site.

**Supplementary figure 4.** Maximum-likelihood phylogenetic tree of the NS gene segment showing the clade 2.3.4.4b H5N1 virus isolated from a tick collected from the slender-billed gull, alongside the virus detected in the same gull. This unrooted tree includes the closest related strains as well as reference sequences representing European genotypes, all belonging to clade 2.3.4.4b. The virus detected in the tick is shown in red, while the virus detected in the gull is shown in blue. Ultrafast bootstrap support higher than 90% is indicated above the nodes. The scale bar indicates the number of nucleotide substitutions per site.

**Supplementary figure 5.** Maximum-likelihood phylogenetic tree of the PB1 gene segment showing the clade 2.3.4.4b H5N1 virus isolated from the slender-billed gull. This unrooted tree includes the closest related strains as well as reference sequences representing European genotypes, all belonging to clade 2.3.4.4b. the virus detected in the gull is shown in blue. Ultrafast bootstrap support higher than 90% is indicated above the nodes. The scale bar indicates the number of nucleotide substitutions per site.

**Supplementary figure 6.** Maximum-likelihood phylogenetic tree of the PA gene segment showing the clade 2.3.4.4b H5N1 virus isolated from the slender-billed gull. This unrooted tree includes the closest related strains as well as reference sequences representing European genotypes, all belonging to clade 2.3.4.4b. the virus detected in the gull is shown in blue. Ultrafast bootstrap support higher than 90% is indicated above the nodes. The scale bar indicates the number of nucleotide substitutions per site.

**Supplementary figure 7.** Maximum-likelihood phylogenetic tree of the NA gene segment showing the clade 2.3.4.4b H5N1 virus isolated from the slender-billed gull. This unrooted tree includes the closest related strains as well as reference sequences representing European genotypes, all belonging to clade 2.3.4.4b. The virus detected in the gull is shown in blue. Ultrafast bootstrap support higher than 90% is indicated above the nodes. The scale bar indicates the number of nucleotide substitutions per site.

