## Supplementary figures and images for "Detection and isolation of H5N1 clade 2.3.4.4b high pathogenicity avian influenza virus from ticks (*Ornithodoros maritimus*) recovered from a naturally infected slender-billed gull (*Chroicocephalus genei*)"

### Supplemental Figure 1

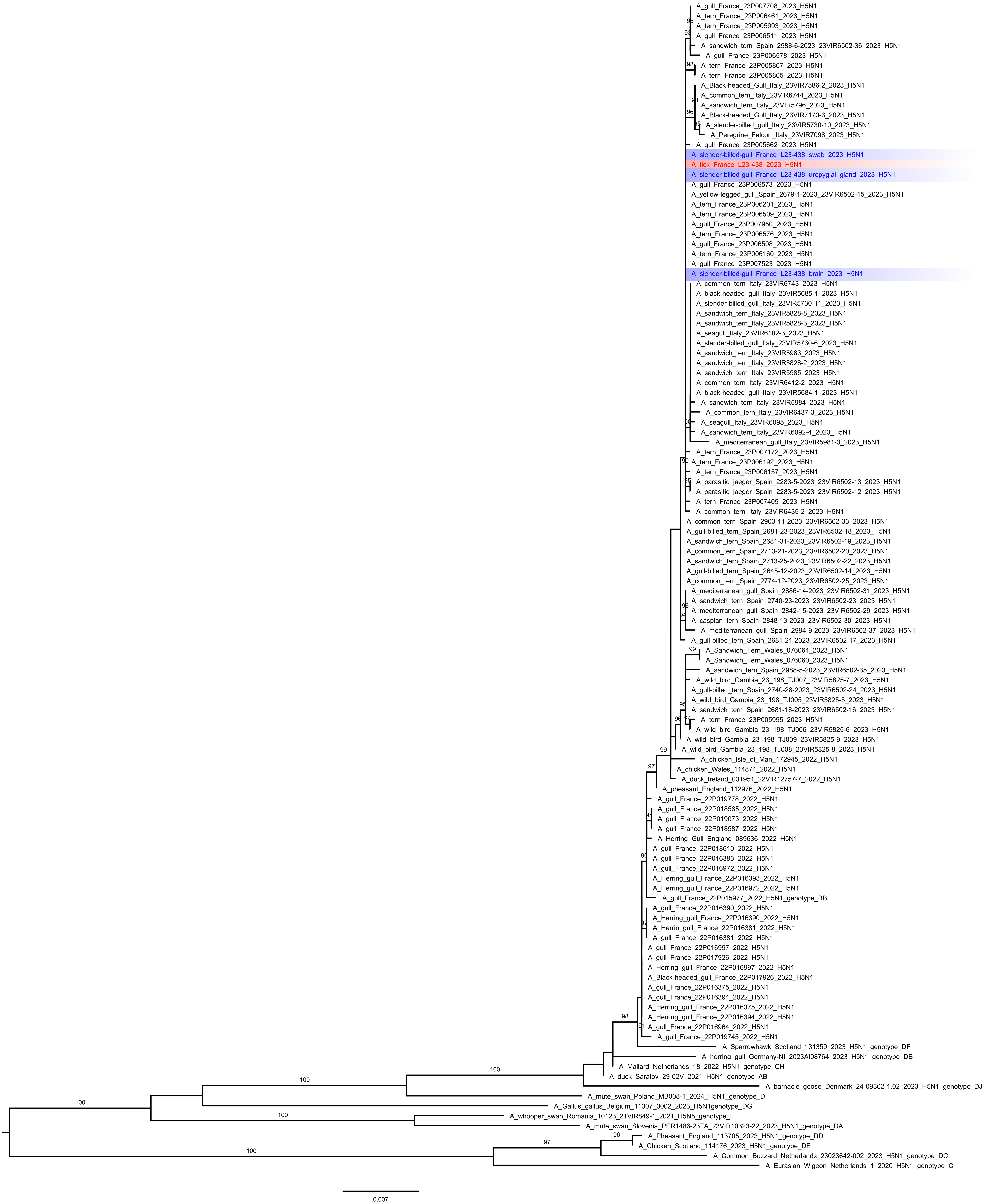

### Supplemental Figure 2

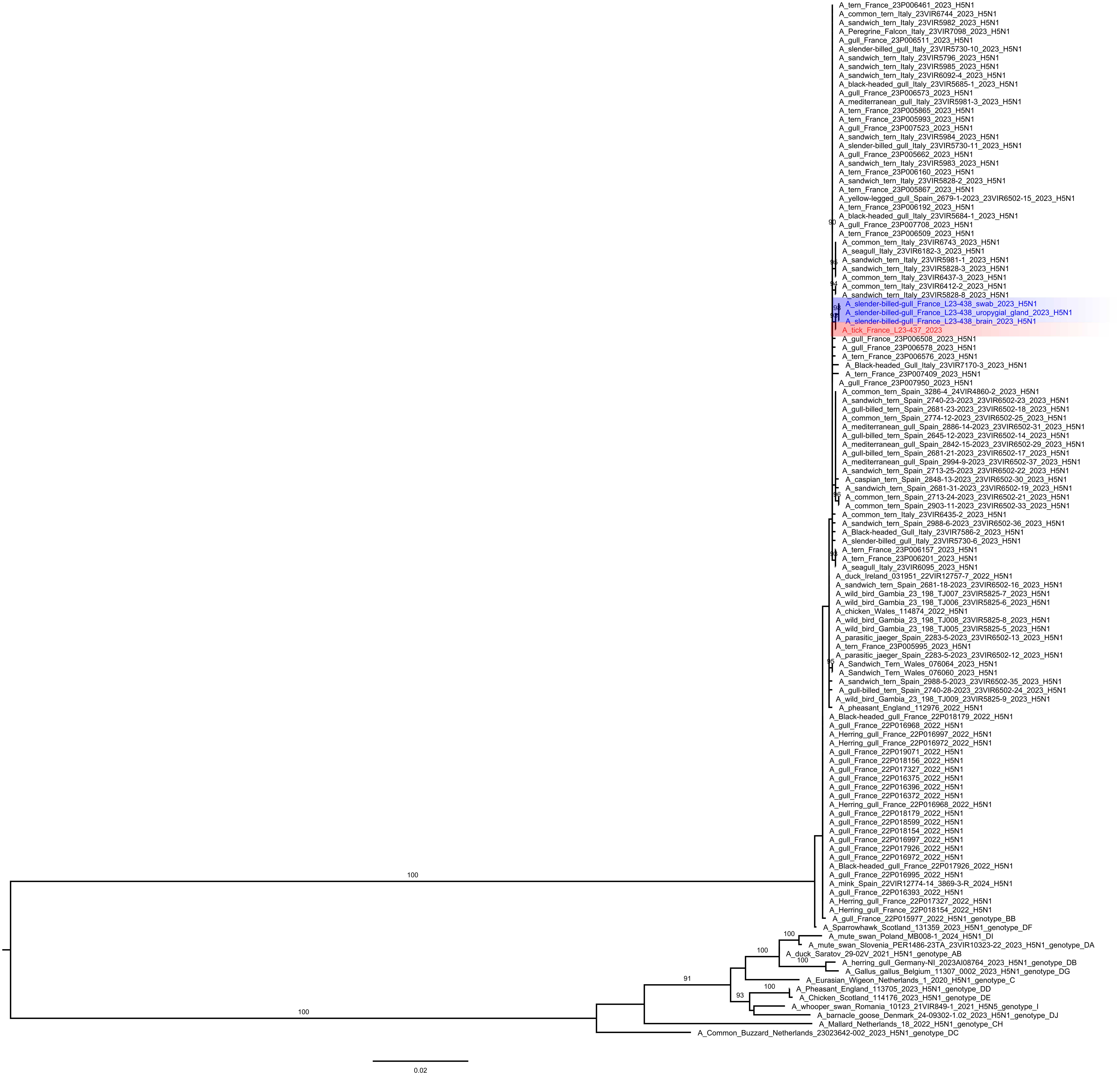

### Supplemental Figure 3

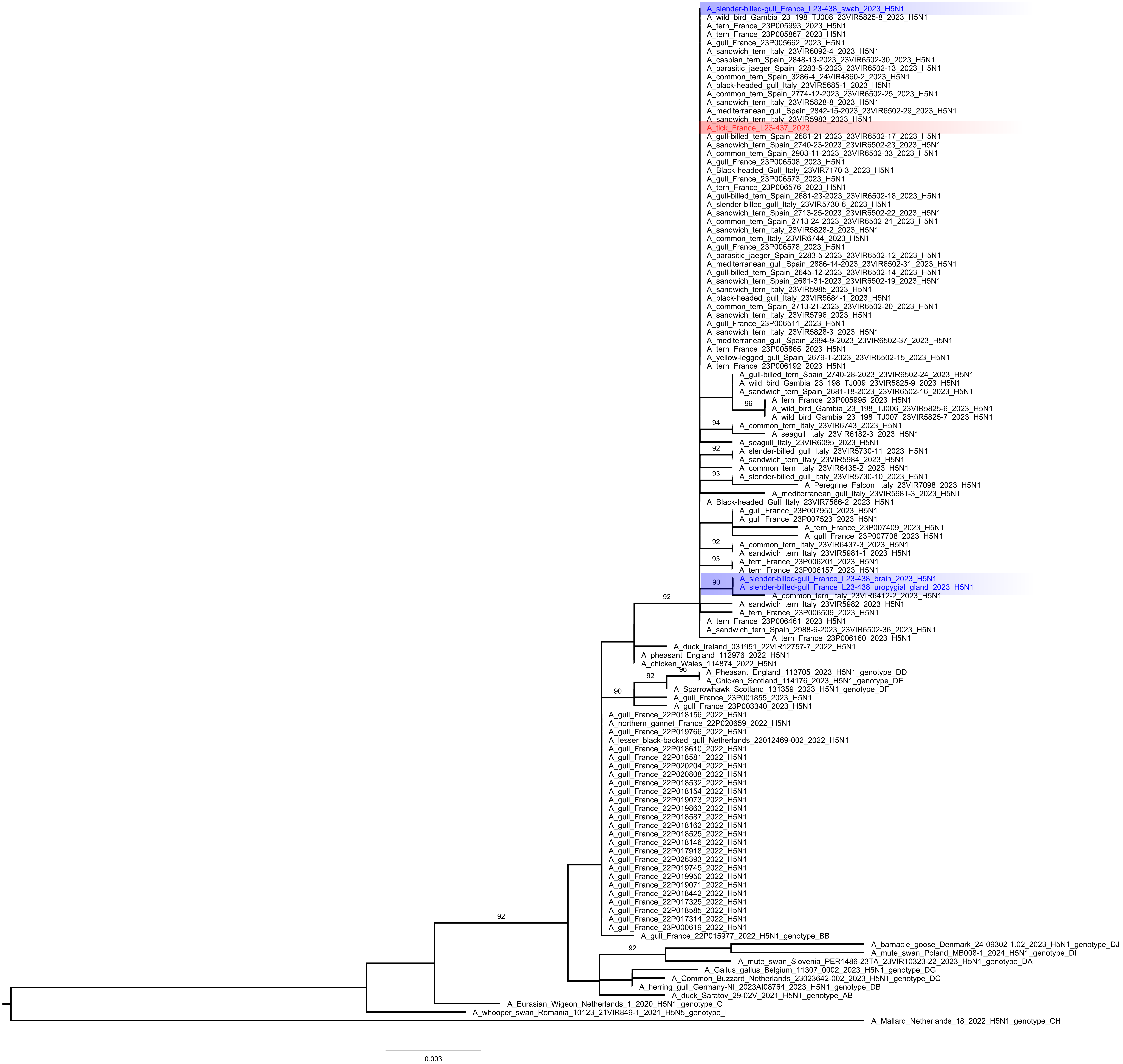

### Supplemental Figure 4

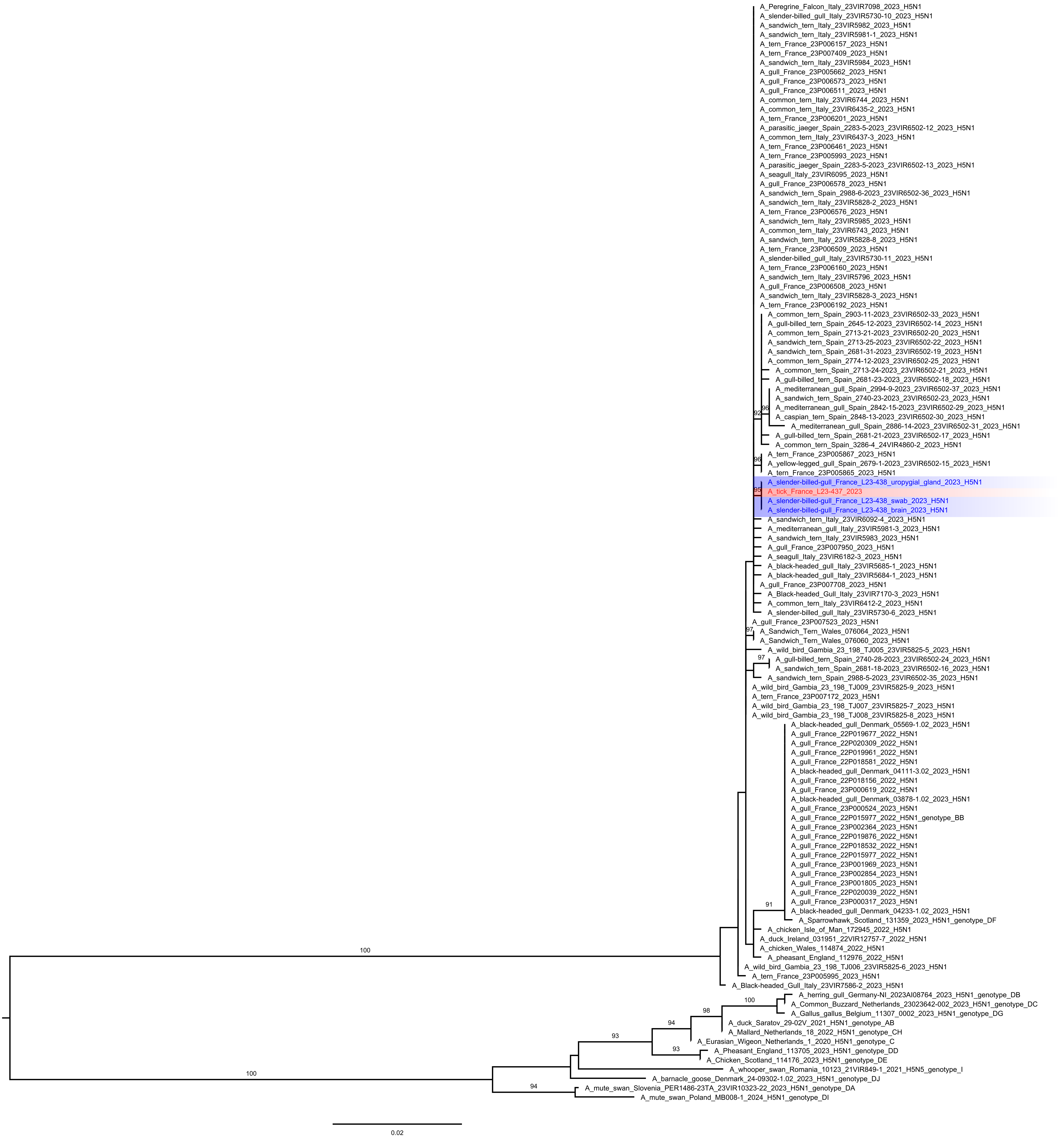

### Supplemental Figure 5

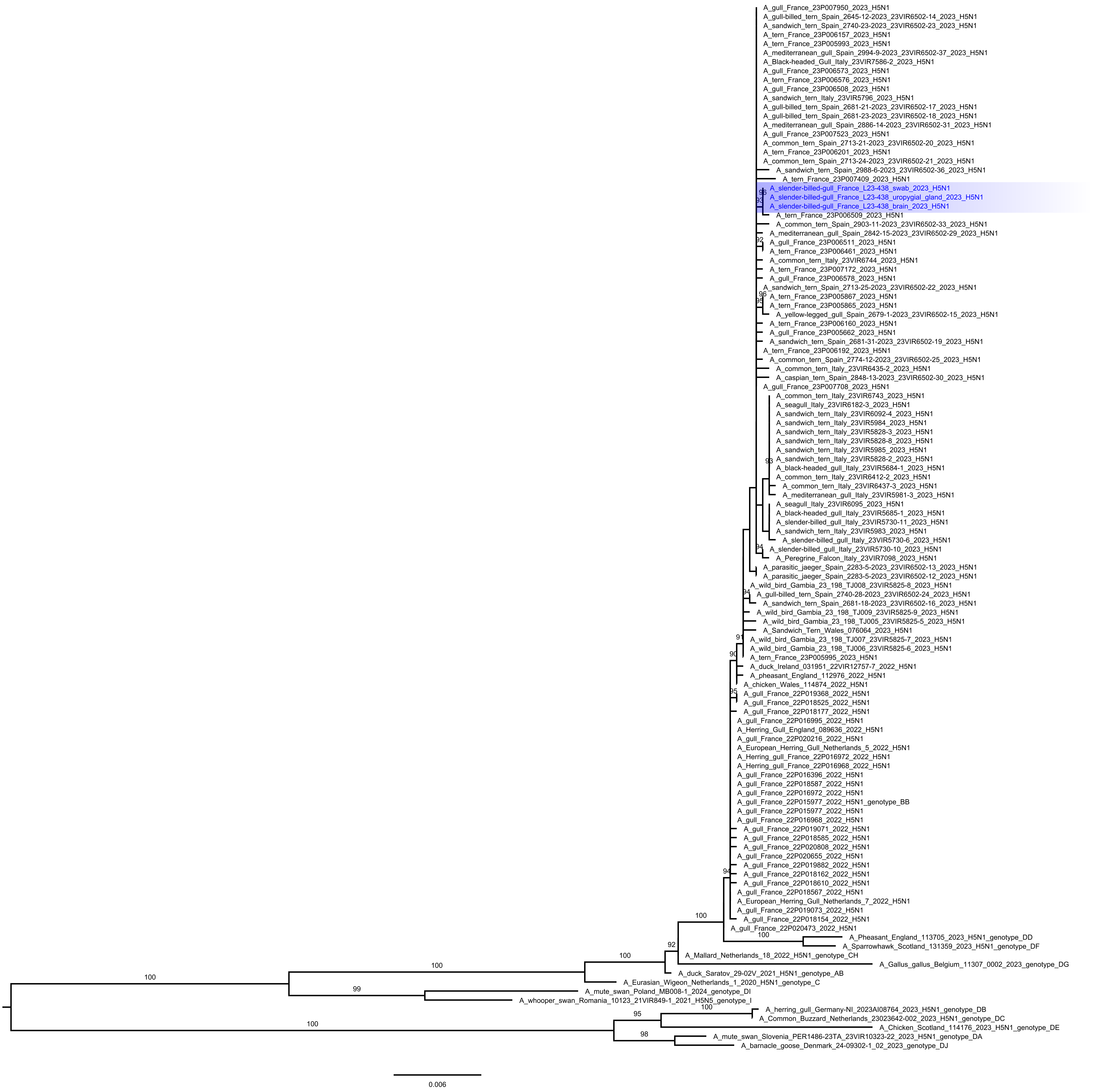

### Supplemental Figure 6

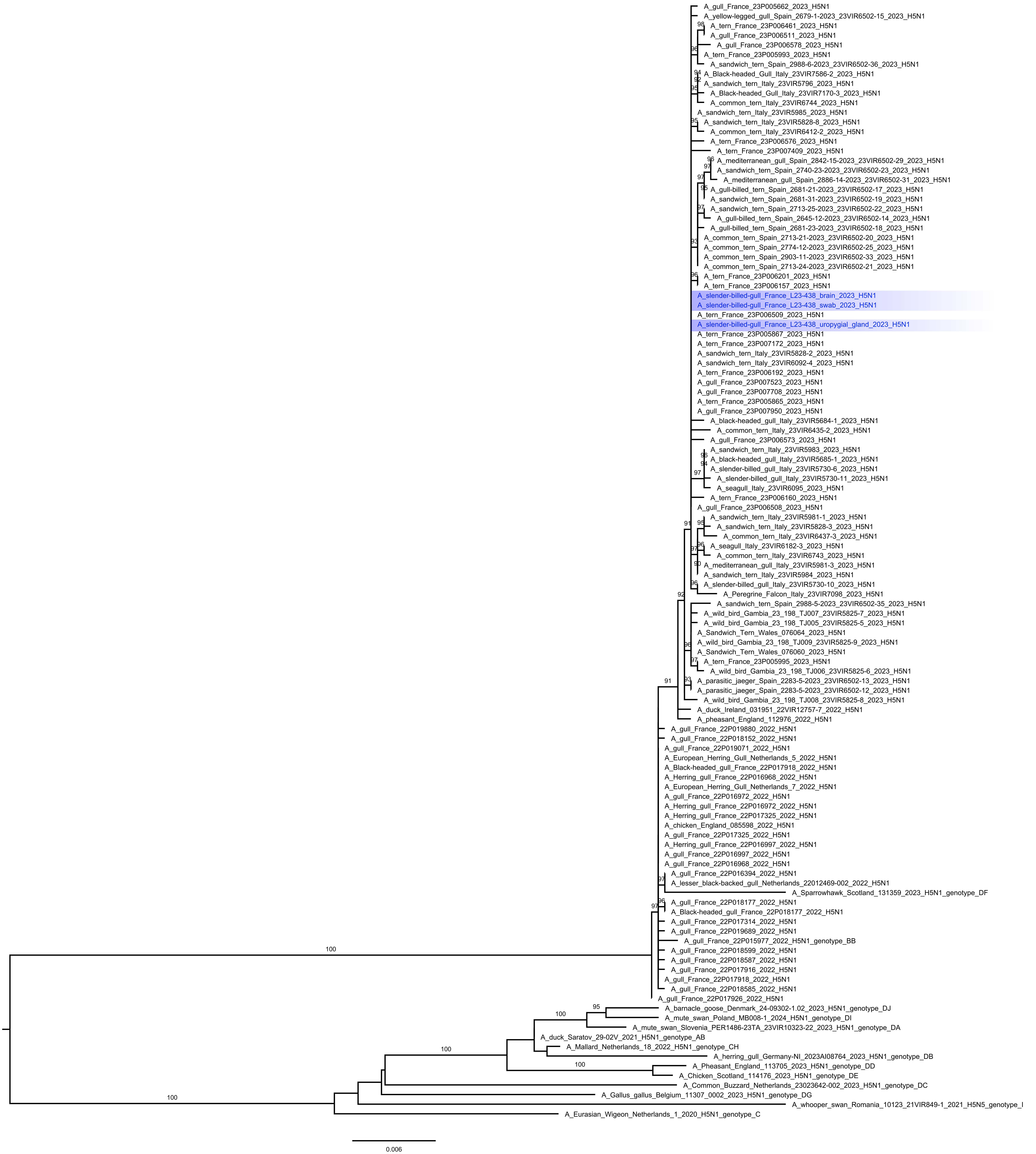

### Supplemental Figure 7

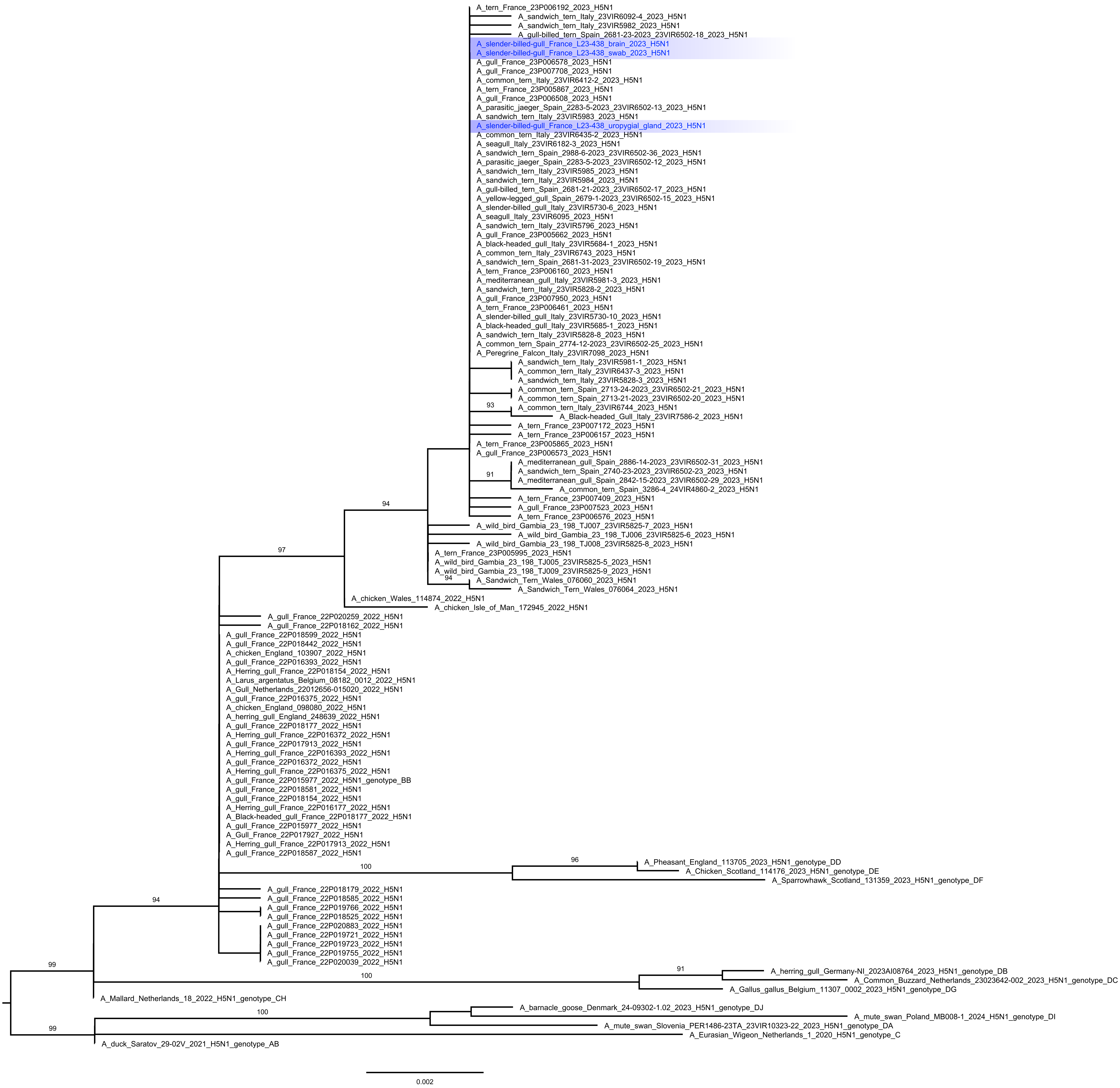
